# Activation of locus coeruleus noradrenergic neurons rapidly drives homeostatic sleep pressure

**DOI:** 10.1101/2024.02.29.582852

**Authors:** Daniel Silverman, Changwan Chen, Shuang Chang, Lillie Bui, Yufan Zhang, Rishi Raghavan, Anna Jiang, Dana Darmohray, Jiao Sima, Xinlu Ding, Bing Li, Chenyan Ma, Yang Dan

## Abstract

Homeostatic sleep regulation is essential for optimizing the amount and timing of sleep, but the underlying mechanism remains unclear. Optogenetic activation of locus coeruleus noradrenergic neurons immediately increased sleep propensity following transient wakefulness. Fiber photometry showed that repeated optogenetic or sensory stimulation caused rapid declines of locus coeruleus calcium activity and noradrenaline release. This suggests that functional fatigue of noradrenergic neurons, which reduces their wake-promoting capacity, contributes to sleep pressure.

## Main

Sleep, an innate behavior essential for our survival^1,2^, is known to be regulated homeostatically: The propensity to sleep and difficulty of awakening (sleep pressure) increase with the amount of prior wakefulness^3^. Several mechanisms of sleep pressure have been proposed^4,5^, including wake-dependent elevation of extracellular adenosine, which can inhibit wake-promoting neurons^6–8^; synaptic strengthening, which can enhance slow-wave synchrony during NREM sleep^9^; and phosphorylation of the proteome and alteration of the transcriptome in the brain^10–13^. In this study, we explore the possibility that activity-induced functional fatigue of wake-promoting neurons contributes to homeostatic sleep pressure. We focused on locus coeruleus (LC) neurons that release noradrenaline/norepinephrine (NE), which strongly promote wakefulness and arousal^14–16^ and are much more active during wakefulness than sleep^17–19^. Using optogenetic and sensory stimulation combined with fiber photometry, we characterized evoked calcium responses and norepinephrine release from LC-NE neurons as well as their effects on sleep-wake states.

To measure the effects of activating LC-NE neurons on sleep-wake brain states, we injected a Cre-inducible AAV expressing a red-shifted channelrhodopsin (AAV2-hSyn-FLEX-ReaChR-mCitrine) bilaterally into the LC of *Dbh-Cre* mice (Fig. 1a). Laser stimulation (10 mW, 532 or 589 nm, 10-ms pulses at 10 Hz or a 2-s pulse every 10 ± 5 s) was applied in 2-min episodes, repeated every 10 min, and sleep-wake states were classified based on electroencephalogram (EEG) and electromyogram (EMG) recordings (Fig. 1b). We found a strong increase in wakefulness within a few seconds after the onset of each stimulation episode (Fig. 1c), consistent with the known wake-promoting effect of LC neurons^14–16^. However, the increase in wakefulness was highly transient: Even within the 2-min stimulation episode, the probability of wakefulness declined to nearly the baseline level, and immediately after the stimulation, wakefulness fell below the baseline, accompanied by increases in NREM and REM sleep. Similar effects were observed with unilateral laser stimulation (Extended Data Fig. 1a) and across different stimulation patterns within each episode (Extended Data Fig. 1b,c).

**Fig. 1.**
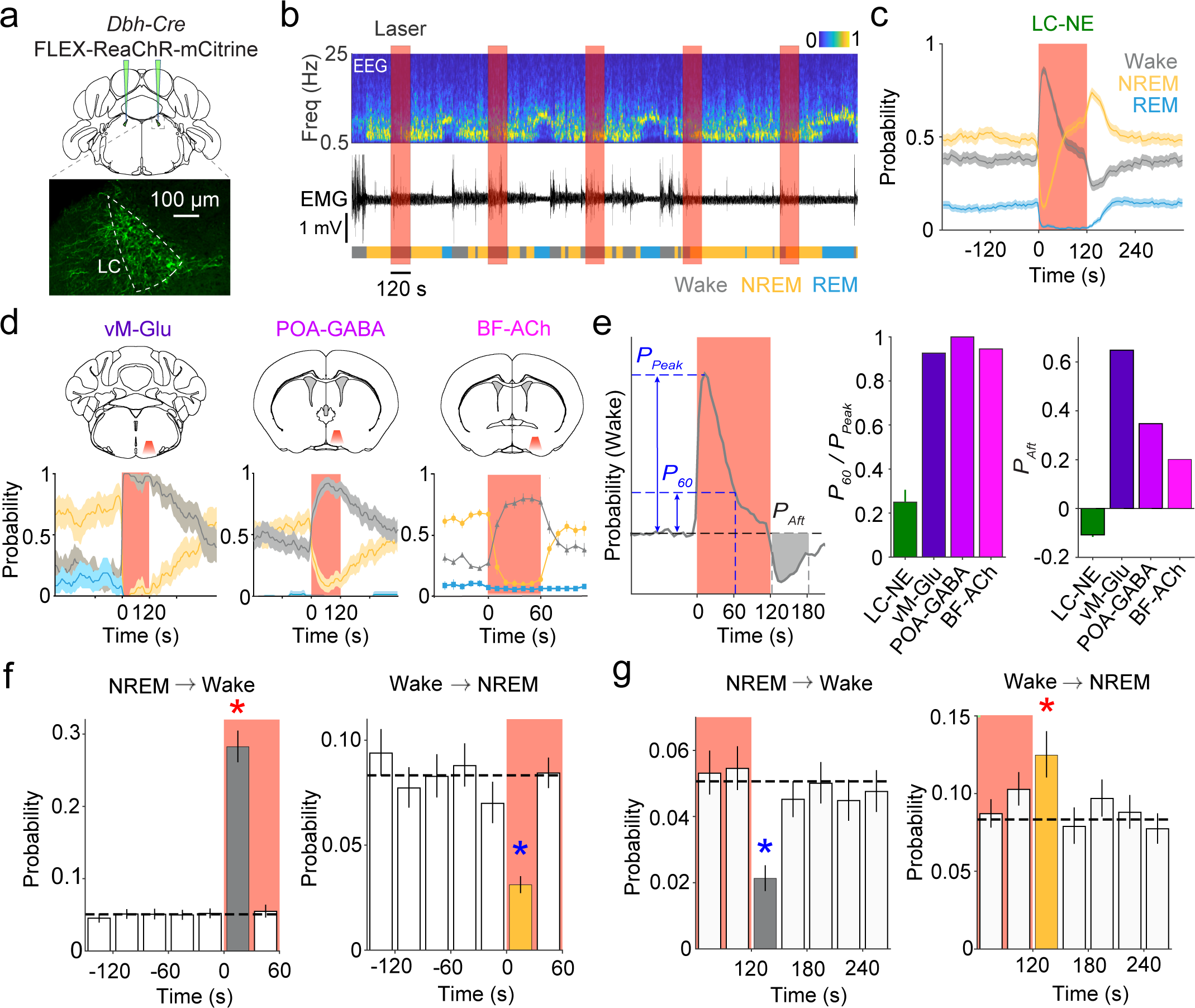
Optogenetic activation of LC-NE neurons drives rebound sleep following transient wakefulness. **a**, Schematic for optogenetic activation of LC-NE neurons. **b**, Example session showing EEG power spectrum (top), EMG (middle) and color-coded brain states (bottom). **c**, Probability of wake, NREM and REM states before, during, and after laser stimulation (mean and 95% confidence interval, n = 8 mice). Red shading, laser stimulation period. **d**, Similar to **c**, but for stimulating glutamatergic neurons in the ventral medulla (vM-Glu), GABAergic neurons in the preoptic area (POA-GABA), and cholinergic neurons in the basal forebrain (BF-ACh); Data reproduced from refs. 20–22. **e**, Comparison of wake-promoting effects of the four populations. Left, schematic illustrating parameters for quantification. Center, persistence of wake effect measured by *P_60_* /*P_peak_* ratio. Right, aftereffects of optogenetic activation (*P_Aft_*). **f**, NREM→wake and wake→NREM transition probabilities before and after stimulation onset. **g**, similar to **f**, but before and after termination of stimulation. Dashed line, mean of baseline period (-240 to 0 s). Red star, significant increase, *p* < 0.0001; Blue star, significant decrease, *p* < 0.0001 (bootstrap).

Such a transient wake-promoting effect of LC neuron activation is in stark contrast to the effects of optogenetic stimulation of glutamatergic neurons in the ventral medulla (vM-Glu)^20^, GABAergic neurons in the preoptic area (POA-GABA)^21^, and cholinergic neurons in the basal forebrain (BF-ACh)^22^, all of which induced wakefulness that outlasted the duration of laser stimulation (Fig. 1d). To quantify the persistence of the wake effect induced by optogenetic activation, we computed the ratio between the increase in wake probability at 60 s after laser onset (*P_60_*) and the peak increase within the laser episode (*P_Peak_*) (Fig. 1e, left). The *P_60_/P_Peak_* ratio for LC-NE neurons was 0.25 ± 0.06 (mean ± s.e.m., n = 8 mice), indicating a strong decay in the wake effect at 60 s after stimulation onset. In contrast, *P_60_/P_Peak_* for vM-Glu, POA-GABA, and BF-ACh neurons were all > 0.9 (Fig. 1e, center). To quantify the aftereffect of the stimulation episode, we computed the difference between the wake probability 0 - 60 s after the stimulation and the baseline period before stimulation (*P_Aft_*). Whereas *P_Aft_* for vM-Glu, POA-GABA, and BF-ACh neurons were all > 0, indicating persistence of the wake effect beyond the stimulation episode, *P_Aft_* for LC-NE neurons was -0.11 ± 0.01, indicative of rebound sleep after the stimulation (Fig. 1e, right).

To further quantify the effects of LC-NE neuron activation, we calculated the probabilities of NREM→wake and wake→NREM transitions before, during, and after the stimulation episode. Compared to the baseline period before stimulation, laser stimulation increased the NREM→wake transition and decreased the wake→NREM transition, but only within the first 30 s of the 2-min episode (Fig. 1f, *p* < 0.0001, bootstrap). Immediately after stimulation, the NREM→wake transition decreased and wake→NREM transition increased significantly (Fig. 1g, *p* < 0.0001), indicating increased sleep drive. In addition to the probabilities of sleep-wake states and their transitions, we also analyzed EEG delta power during NREM sleep, a common measure of homeostatic sleep pressure. After a transient decrease at the onset of stimulation, NREM delta power quickly rebounded and rose above the baseline level immediately after stimulation (Extended Data Fig. 1d). Together, the increases in both NREM probability and EEG delta power indicate an elevation of sleep pressure after LC neuron activation.

A potential mechanism for the rapid decrease in wake-promoting effect of LC-NE neurons is an attenuation of their activity caused by repeated optogenetic activation. We next measured calcium activity of LC-NE neurons in response to laser stimulation. Cre-inducible AAVs expressing jGCaMP8m (AAV9-hSyn-FLEX-jGCaMP8m^23^) and ReaChR (AAV2-hSyn-FLEX-ReaChR-mCitrine) were injected into the LC of *Dbh-Cre* mice, and optogenetic stimulation and fiber photometry imaging were performed simultaneously through the same optic fiber (Fig. 2a). Without optogenetic stimulation, LC-NE neurons showed higher activity during wakefulness than both NREM and REM sleep (Extended Data Fig. 2), consistent with previous studies^17–19^. Each 2-s laser pulse evoked a salient calcium response (Fig. 2b), but the response amplitude decreased consistently over successive laser pulses within each episode (Fig. 2c). For pulse 12 (near the end of each episode), the response amplitude was only 59% ± 3% (s.e.m.) of the response to pulse 1 (Fig. 2d).

**Fig. 2.**
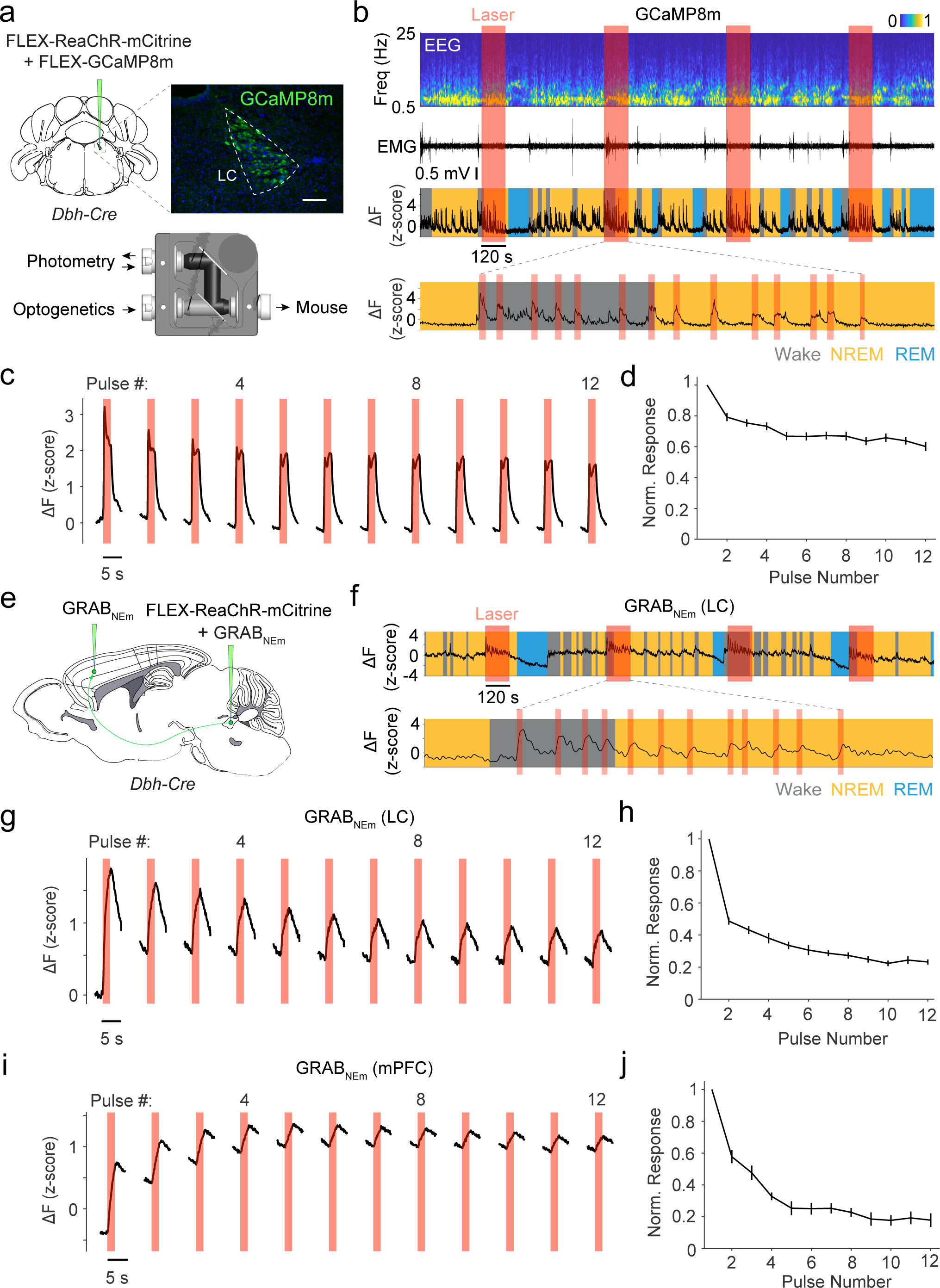
Calcium activity of LC-NE neurons and NE release evoked by optogenetic activation. **a**, Top, coronal diagram for injection of Cre-inducible ReaChR-mCitrine and GCaMP8m into the LC of *Dbh-Cre* mice and fluorescence image showing GCaMP8m expression (immunostaining). Scale bar, 100 µm. Bottom, optical pathway for simultaneous optogenetic stimulation and fiber photometry. **b**, Example session showing calcium activity in LC-NE neurons during optogenetic stimulation. Expanded view of one stimulation episode shows calcium responses to individual 2-s pulses. **c**, Calcium responses to individual 2-s pulses averaged across all episodes and all mice (n = 19 sessions recorded from 6 mice). **d**, Summary of response amplitude (normalized by the maximal response in each episode, mean ± s.e.m.). **e**, Sagittal diagram for injection of ReaChR-mCitrine and GRAB_NEm_ into the LC and GRAB_NEm_ into the mPFC of *Dbh-Cre* mice. **f**, NE release in the LC evoked by optogenetic activation. **g-j**, Similar to **c,d**, but for NE responses in the LC (**g,h**; n = 16 sessions, 7 mice) and mPFC (**i,j**; n = 10 sessions, 6 mice).

We next measured laser-evoked NE release by expressing a GPCR-activation-based fluorescent NE sensor (GRAB_NEm_)^24^ in the LC or the medial prefrontal cortex (mPFC) and ReaChR in LC-NE neurons (Fig. 2e). Each 2-s laser pulse evoked a rapid increase in NE level (Fig. 2f), but the response amplitude also declined progressively in both the LC (Fig. 2g,h) and the mPFC (Fig. 2i,j). In fact, the decrease in evoked NE release measured by the ratio between pulse 12 and pulse 1 (LC: 0.23 ± 0.02; mPFC, 0.18 ± 0.04; mean ± s.e.m.) was much greater than the decrease in calcium responses (Fig. 3j), indicating a stronger decline of the functional output of these neurons. Thus, repeated LC neuron activation caused a rapid “functional fatigue” – evidenced by diminished calcium activity and NE release – which may contribute to the loss of their wake-promoting effect within each stimulation episode (Fig. 1f). The sleep rebound immediately after stimulation (Fig. 1c,g) is likely caused by a reduction of spontaneous LC-NE activity following the 2-min optogenetic activation.

**Fig. 3.**
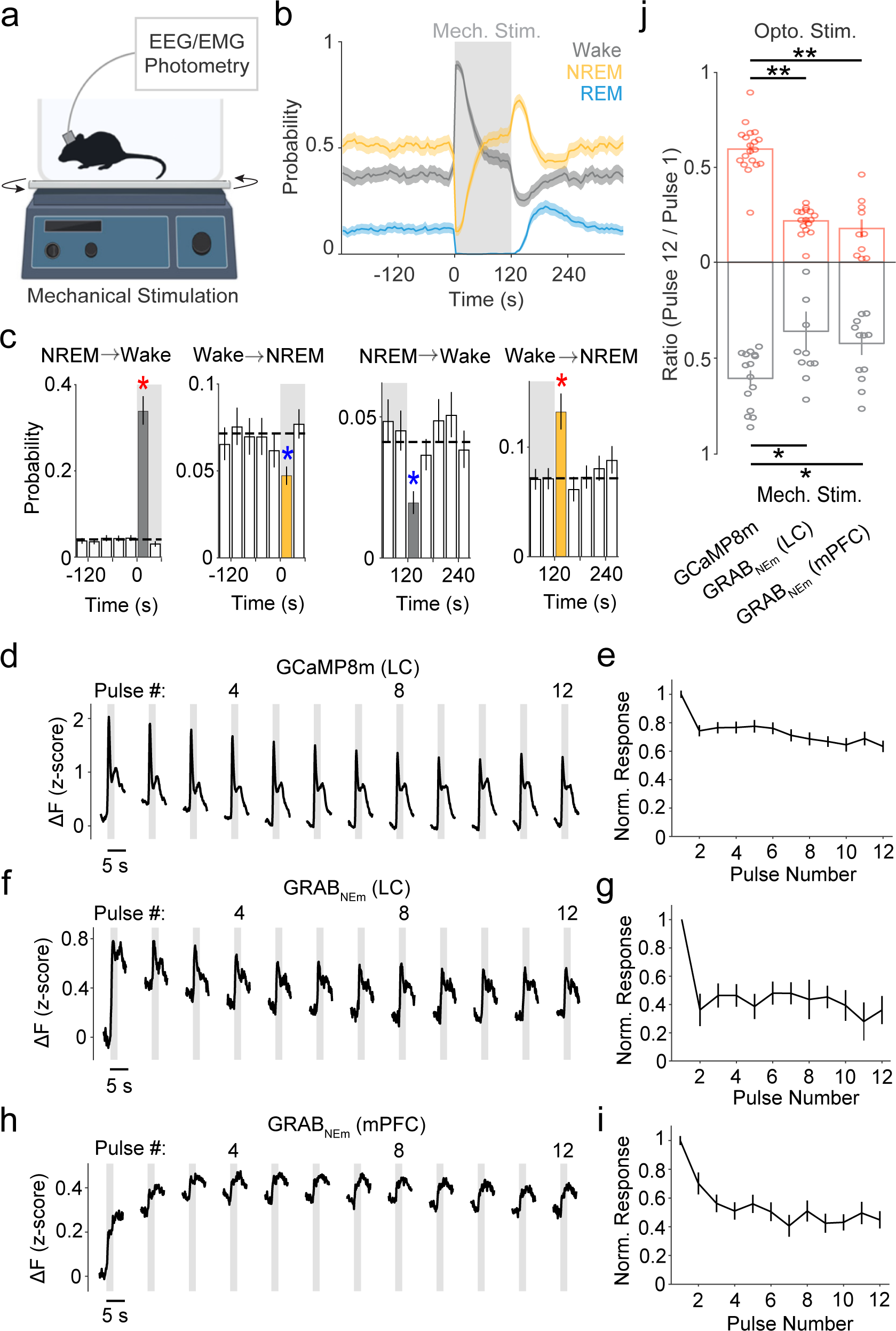
Effects of sensory stimulation on brain states, LC neuron calcium activity and NE release. **a**, Schematic of mechanical stimulation with an orbital shaker. **b**, Sleep-wake probability before, during, and after 2-min mechanical stimulation (mean and 95% confidence interval, n = 6 mice). **c**, NREM→wake and wake→NREM transition probabilities immediately before and after the onset (left) and termination (right) of stimulation. Red star, significant increase, *p* < 0.0001; Blue star, significant decrease, *p* < 0.0001 (Bootstrap). **d,e**, Calcium responses of LC-NE neurons to consecutive pulses of mechanical stimulation (n = 15 sessions, 5 mice), similar to Fig. 2c**,d**. **f-i**, Similar to **d,e**, but for mechanical stimulation-evoked NE release in the LC (**f,g**; n = 11 sessions, 5 mice) and mPFC (**h,i**; n = 13 sessions, 6 mice). **j**, Summary of functional fatigue induced by optogenetic (top) and sensory stimulation (bottom), quantified by Pulse 12/Pulse 1 ratio for calcium responses and NE release (mean ± s.e.m.); *, *p* < 0.01 **, *p* < 0.0001 (one-sided, Wilcoxon rank-sum test).

Since optogenetic stimulation could potentially induce non-physiological patterns of neuronal activity, we wondered whether repeated sensory stimulation could also induce LC neuron fatigue and homeostatic sleep pressure. The mouse was recorded in its home cage on top of an orbital shaker (Fig. 3a), which provided mechanical stimulation (cage shaking) in the same temporal pattern as optogenetic stimulation (2-s stimulation every 10 ± 5 s, for 2 min/episode). Such mechanical stimulation also induced a transient wakefulness followed by a sleep rebound (Fig. 3b,c), with a time course similar to that induced by optogenetic stimulation of LC-NE neurons (Fig. 1c,f,g). Furthermore, calcium activity of LC-NE neurons (Fig. 3d,e) and NE release in the LC (Fig. 3f,g) and mPFC (Fig. 3h,i) evoked by each 2-s mechanical stimulation decreased progressively over the 2-min episode, and the decrease in NE release was significantly stronger than the decrease in calcium responses (Fig. 3j). Thus, repeated sensory stimulation can also induce functional fatigue of LC-NE neurons as well as homeostatic sleep pressure, similar to the effects of optogenetic stimulation. When optogenetic and sensory stimulation episodes were applied consecutively (Extended Data Fig. 3a-c), the wake-promoting effect of sensory stimulation was significantly reduced by the preceding optogenetic stimulation (Extended Data Fig. 3d); similarly, the wake-promoting effect of optogenetic stimulation was significantly reduced by the preceding sensory stimulation (Extended Data Fig. 3f). This is consistent with our finding that repeated sensory and optogenetic stimulation both lead to functional fatigue of LC-NE neurons, which can reduce the wake-promoting effect of succeeding stimulation.

In sum, we have shown that activation of LC-NE neurons with either optogenetic or sensory stimulation caused rapid declines of their calcium responses and evoked NE release (Figs. 2,3), which may contribute to the increase in sleep propensity (Fig. 1) and sleep depth (Extended Data Fig. 1d). The highly transient increase in wakefulness, which distinguishes LC-NE neurons from other wake-promoting neurons (Fig. 1d,e), is consistent with previous reports that activation of LC-NE neurons often drives brief wakefulness and microarousals^16^. The cellular processes mediating the rapid decline of LC neuron calcium responses and NE release remain unclear. A possible mechanism is autoinhibition induced by released NE, which binds to α_2_ adrenergic receptors and causes hyperpolarization of LC neurons following their activation^25,26^. In addition to direct effects on LC neurons, released NE can also affect many other neurons, astrocytes, and microglia which, for example, may lead to increased adenosine that could inhibit LC-NE neurons^27^ and increase sleep pressure^6–8^. It would be interesting to explore how manipulations of these pathways may impact activity-induced decline of LC-NE activity and homeostatic sleep regulation.

In general, sleep pressure could be driven by an increase in sleep-promoting factors (e.g., adenosine) and/or weakening of wake-promoting mechanisms, such as functional fatigue of LC-NE neurons. LC-NE neurons are known to be particularly vulnerable to neurodegeneration, exhibiting earlier and greater cell loss than many other neurons in both Alzheimer’s and Parkinson’s diseases^28–30^. An intriguing possibility is that their susceptibility to functional fatigue is mechanistically linked to their vulnerability to degeneration. The rapid decline of LC-NE functional efficacy could reflect sensitivity of these neurons to activity-induced metabolic stress (e.g., oxidative stress caused by reactive oxygen species)^31^ that may contribute to both functional fatigue and neurodegeneration^32,33^. On the other hand, the resulting increase in sleep pressure could offer a protective mechanism against the harmful effects of excessive LC-NE activation. Thus, the dynamic regulation of LC-NE activity may represent a key link in the bi-directional relationship between sleep impairment and neurodegeneration^34–36^.

## Methods

### Animals

All procedures were performed in accordance with the protocol approved by the Animal Care and Use Committee at the University of California, Berkeley. Adult (6-12 weeks old) male and female *Dbh-Cre* mice (B6.FVB(Cg)-Tg(Dbh-cre)KH212Gsat/Mmucd, MMRRC: 036778-UCD) were used for all experiments. Mice were kept on a 12:12 light:dark cycle. After virus injections and surgical implantation of EEG/EMG electrodes and optical fibers, mice were individually housed to prevent damage to the implant before experiments. Experiments were conducted 2-4 weeks after surgery.

### Surgeries

Anesthesia was induced with 5% isoflurane and maintained with 1.5% isoflurane on a stereotax. Buprenorphine (0.1 mg/kg, SC) and meloxicam (10 mg/kg SC) were injected before surgery. Lidocaine (0.5%, 0.1 mL, SC) was injected near the target incision site. Body temperature was kept stable throughout using a heating pad and a feedback thermistor. After sterilizing the skin with ethanol and betadine, a circular patch of skin and connective tissue was cut away to expose the skull. Surgeries typically consisted of virus injection followed by EEG/EMG implantation and fiber implantation.

For virus injections, a craniotomy was drilled above the target site (see below for details of virus injections). 10 minutes after virus injection, the injection pipette was slowly removed from the injection site. For recording the electroencephalogram (EEG) and electromyogram (EMG), stainless-steel wires were soldered to a 20-pin header (Minitek 127T series). EEG screw electrodes were implanted in both sides of the frontal lobe (± 1.5 mm ML, -1.5 mm AP). EMG stainless-steel wire electrodes (0.003” diameter) were inserted into both sides of the trapezius muscle. Reference screws were inserted into each side of the cerebellum. The EEG/EMG implant was secured by dental cement before fiber implantation. For fiber photometry and optogenetic stimulation, a fiber implant (1.25-mm ferrule, 200-µm core) was held by a stereotactic fiber holder (XCL, Thor Labs). Either single (Neurophotometrics) or dual fibers (TFC_200/245-0.37_6mm_TS1.8_FLT, Doric Lenses Inc.) were implanted 100 µm above the virus injection site. Fibers were secured with cyanoacrylate glue and dental cement before withdrawing the stereotactic fiber holder.

### Virus Injections

Injections were performed using a Nanoject II (Drummond Scientific) with glass pipettes. The injection settings were set to a 50 nL injection volume at a rate of 23 nL/s with a 30-s interval between injections. AAV2-hSyn-FLEX-ReaChR-mCitrine (0.3-3.0 × 10^13^ gc/mL, 50955 from Addgene) was injected bilaterally or unilaterally into the LC of *Dbh-Cre* mice (-5.5 AP, ±0.9 ML, 250 nL at -3.7 DV and 250 nL at -3.5 DV). For fiber photometry experiments, the ReaChR virus was co-injected unilaterally into the LC with AAV9-hSyn-FLEX-jGCaMP8m (3 × 10^13^ gc/mL, 162378-AAV9 from Addgene) or with AAV9-hSyn-GRAB_NEm_ (2-9 × 10^13^ gc/mL, WZ Biosciences). For photometry in the mPFC, AAV9-hSyn-GRAB_NEm_ was injected unilaterally (250 nL, +2.1 AP, +0.3 ML, -1.6 DV).

### EEG/EMG recording

EEG/EMG electrodes were connected to a custom recording cable (P1 Technologies) that interfaced with data acquisition hardware (S-Box-16, PZ-5, RZ5 from Tucker-Davis Technologies). Mice were typically habituated overnight and recording sessions started at 6:30 am and lasted 6 hours. Data was acquired using Synapse software (version 95-44132P, Tucker-Davis Technologies), with a digital bandpass filter of 1-750 Hz and sampling rate of 1500 Hz. Classification of sleep-wake behavioral state in each 5-s epoch was performed according to our published machine-learning method and visual inspection using established criteria and a graphical user interface in Matlab^20,22,37^. Briefly, NREM sleep was characterized by low muscle tone (root-mean-square of EMG voltage) and high EEG delta power (sum of power from 1-5 Hz). REM sleep was characterized by low delta power, high theta power (sum of power from 6-9 Hz) / delta power ratio, and low muscle tone. Wakefulness included active wake with high muscle tone and quiet wake with low delta power and low muscle tone with low theta/delta power ratio.

### Optogenetic stimulation

To study the transient wakefulness induced by activation of LC-NE neurons, we applied laser stimulation (532 nm or 589 nm, 10-mW at fiber tip). Recording sessions consisted of 24 stimulation episodes (first stimulation typically at 8:00 am, inter-stimulation-episode-onset-interval of 10 min). Each stimulation episode lasted 2 minutes and consisted of either phasic (2-s pulse, 10 ± 5 s inter-stimulation-onset-interval) or tonic (10-ms pulse, 10-Hz) stimulation protocols controlled by TTL pulses from an RZ5 BioAmp Procesor (Tucker-Davis Technologies). Three 6-hour sessions were typically recorded from each mouse.

### Fiber photometry

Fiber photometry was performed using a CMOS-camera-based system (FP3001, Neurophotometrics), with 470 nm and 405 nm LEDs. The sampling rate for each channel was 10 frames/s. The system was turned on for 70 minutes to equilibrate the camera and bleach autofluorescence prior to collecting experimental data. For analysis of jGCaMP8m and GRAB_NEm_ signals, we used a ratiometric method by first subtracting the autofluorescence of a fiber implant (recorded from an isolated fiber in a dark environment for 6 hours) and then calculating the ratio between signals in the 470 nm and 405 nm channels. The result was then z-scored. For the example trace in Fig. 2f, the photometry signal was low-pass filtered at 0.2 Hz.

### Mechanical stimulation

An Internet-of-Things relay device (Digital Loggers) was used to control motion (∼60 revolutions per minute) of an orbital shaker (MT-201-BD, Labnique) with a TTL pulse from an RZ5 BioAmp Processor (Tucker-Davis Technologies). The home cage of the mouse was placed on a sheet of aluminum foil grounded to the recording equipment through the PZ-5 grounding port (Tucker-Davis Technologies). Mechanical stimulation often introduced sharp artifacts in the photometry data that were clearly distinguishable from the relatively slow calcium and NE fluctuations. To remove these artifacts, we calculated the derivative of the fluorescence signals and removed data points that rose above a threshold value.

### Quantification and normalization of calcium and NE responses

For each optogenetic or mechanical stimulation pulse, the response was quantified by calculating the mean signal from 0.2 to 0.6 s after stimulation onset minus the mean signal from -1.5 to 0 s before stimulation onset. This short response window was chosen because the mechanical stimulation experiments involved 2 revolutions of the orbital shaker within the 2 s pulse (60 revolutions/min), which often produced two peaks in the calcium and NE responses, whereas the optogenetic stimulation was a single square pulse which typically evoked a single peak (within the 0.2 to 0.6 s window for calcium responses). The response for each pulse in the episode was averaged across all stimulation episodes in a recording session. For normalization, the average response for each pulse was divided by the maximum average response from the session.

### Statistics

Sleep-wake probability plots and transition probabilities were analyzed as described previously^20^. Briefly, the 95% confidence intervals (CIs) for brain state probabilities and statistical tests were calculated from bootstrapped data as follows: For n mice, we calculated the CI by randomly resampling the data (with replacement) for 10,000 iterations. Then, we recalculated the mean probabilities for each sleep-wake state across the mouse averages for the n mice. The lower and upper CIs were then extracted from the distribution of the bootstrapped data. To test whether a given transition probability was significantly larger or smaller from the baseline at stimulation onset or at recovery onset, we performed bootstrap significance tests comparing the target bin and the baseline average. The photometry data was averaged across sessions to minimize noise from certain mice that were only recorded in one session. We used pairwise t-tests to compare the downregulation of calcium and NE responses.

### Histology

To evaluate the location and strength of virus expression as well as the fiber position, mice were anesthetized with isoflurane before transcardial perfusion with 15 mL DPBS (14200-075, Gibco) and 30 mL 4% paraformaldehyde (15714, Electron Microscopy Sciences) in DPBS. For fiber localization, the bottom of the skull was removed to expose the ventral surface of the brain and the brain was post-fixed in 4% paraformaldehyde DPBS for 1-2 nights before separating the brain from the top of the skull. The fixed brain was dehydrated in 15 mL of 30% sucrose DPBS for 1-2 nights, allowing time for it to sink to the bottom of a conical tube. The brain was then embedded in tissue freezing medium (General Data Company, Inc.) and frozen in a -80°C freezer for at least 1 hour. Brains were cryosectioned at 40 µm using a cryostat (Leica). ReaChR-mCitrine fluorescence was visible without immunostaining. Nonspecific binding was blocked with 10% normal donkey serum in PBS-T (pH 7.2, 0.1% Tween-20). To identify the LC region, cryosections were immunostained with a rabbit tyrosine hydroxylase antibody (ab112, abcam, 1:300 in blocking solution). To evaluate jGCaMP8m and GRAB_NEm_ expression, cryosections were immunostained with a chicken GFP antibody (GFP-1020, Aves Labs, 1:100). Brain sections were stained for one night at 4°C. Donkey-anti-rabbit antibodies conjugated to red or far-red Alexa fluorophores (A10042, Invitrogen; ab150067, abcam) were used to visualize tyrosine hydroxylase staining. A donkey-anti-chicken antibody conjugated to Alexa-488 (A78948, Thermo Fisher) was used to visualize GFP staining. Secondary antibodies were incubated for 2 h at room temperature. Slides were washed with PBS-T, mounted with Aqua-Poly/Mount (18606, Polysciences), and imaged on a fluorescence microscope (Keyence).

### Reporting summary

Further information is available in the Nature Portfolio Reporting Summary.

## Data availability

Source data are provided in this paper and all other data related to this work will be provided upon request.

## Code availability

All code used to analyze the presented data will be provided upon request.

## Biological materials availability

All viruses and mouse lines used in this work will be provided upon request.

## Acknowledgments

Funding for this work was from NIH NINDS U01NS113358, the Howard Hughes Medical Institute, and the James R. “Jim Bob” Moffett, Sr. Postdoctoral Fellowship from the Parkinson’s Foundation (DS). We also thank K. Ritola and the UNC BRAIN Initiative Viral Vector Core for viral packaging and S. Velazquez for building sleep-wake recording boxes to expand recording capacity.

## Author Contributions

D.S. and Y.D. designed experiments and wrote the manuscript. D.S. performed all optogenetic and mechanical stimulation experiments with assistance from C.C. Construction of EEG/EMG implants, cyrosectioning, immunostaining and data analyses were carried out by D.S., C.C., S.C., L.B., Y.Z., R.R., and A.J. Control animals with GRAB_NEm_-mutant sensor injected in the LC were provided by D.D. and J.S. Viral constructs and antibodies as well as technical advice on surgical procedures, histology, and data analysis were provided by X.D., B.L., and C.M. All authors provided feedback throughout the project.

## Figure legends

**Extended Data Fig. 1.**
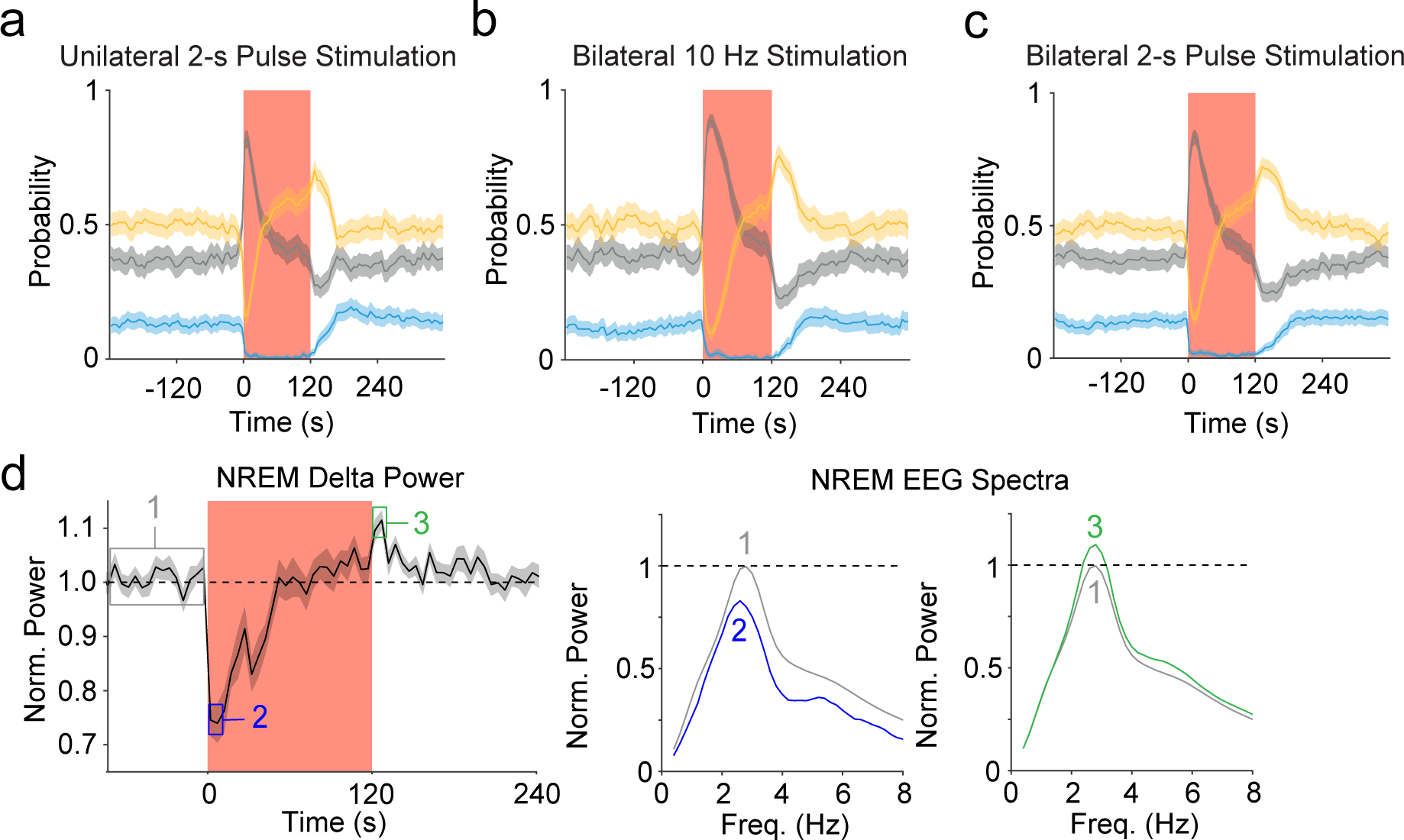
Effects of different patterns of LC-NE neuron optogenetic activation on brain states and NREM EEG. **a**, Probability of wake, NREM and REM states before, during, and after unilateral LC stimulation (2-s pulses every 10 ± 5 s, n = 8 mice). **b,c**, similar to **a**, with bilateral stimulation: 10 ms pulses at 10 Hz, n = 8 mice (**b**) or 2-s pulses every 10 ± 5 s, n = 8 mice (**c**). **d**, Left, EEG delta power before, during, and after LC stimulation, averaged across all sessions (n = 44 sessions, 8 mice). Right, EEG power spectra within three time windows: before stimulation (1, gray box), soon after stimulation onset (2, blue), and immediately after stimulation (3, green), normalized by the peak in window 1.

**Extended Data Fig. 2.**
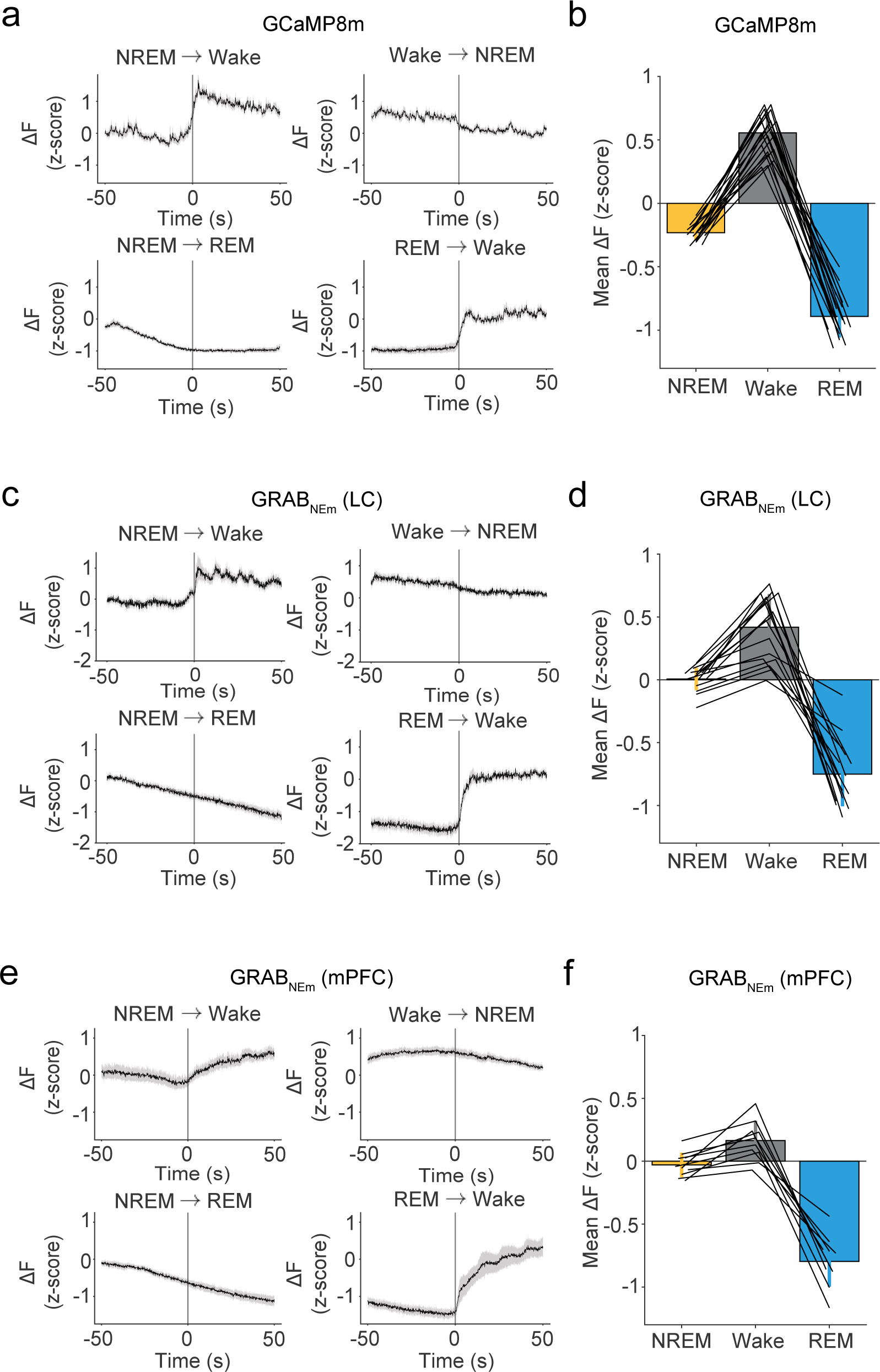
LC-NE neuron activity at sleep-wake state transitions. **a**, Calcium signal in LC-NE neurons before and after each state transition. **b**, Mean calcium signal during each state (mean ± s.e.m.), each line represents one session (n = 19 sessions, 6 mice). **c-f**, Similar to **a,b**, but for NE levels in the LC (**c,d**; n = 16 sessions, 7 mice) and mPFC (**e,f**; n = 10 sessions, 6 mice).

**Extended Data Fig. 3.**
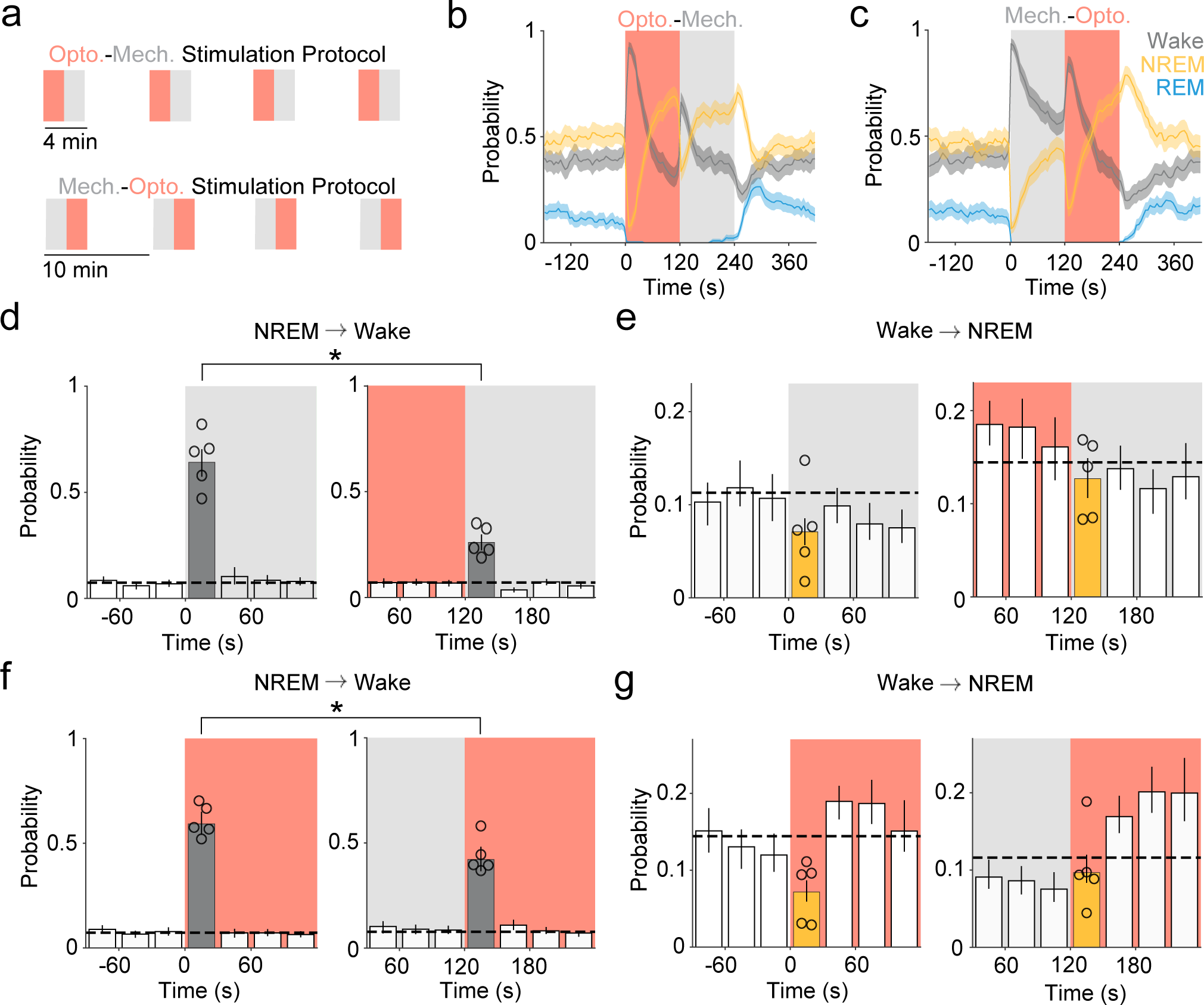
Effects of consecutive optogenetic and mechanical stimulation. **a**, Schematic for consecutive optogenetic and mechanical stimulation episodes. **b,c**, Probability of each state before, during, and after optogenetics-mechanical stimulation (**b**) and mechanical-optogenetic stimulation (**c**). **d**,**e**, NREM→Wake (**d**) and Wake→NREM (**e**) transition probabilities at mechanical stimulation onset with mechanical-optogenetic (left) or optogenetic-mechanical protocol (right). The increase in NREM→Wake probability evoked by mechanical stimulation is significantly suppressed by the preceding optogenetic stimulation. **f**, Similar to **e**, showing that NREM→Wake probability evoked by optogenetic stimulation is significantly suppressed by the preceding mechanical stimulation. *, *p* < 0.05 (Wilcoxon signed-rank test, n = 5 mice, 3 sessions for each protocol).

**Extended Data Table 1.** Test statistics and *p*-values for all figures.

| Extended Data Table 1. Test statistics and *p*-values for all figures. |  |  |  |
| --- | --- | --- | --- |
| Figure | Sample Size | Statistical Test | Results |
| Fig. 1f | 8 mice, 2 stimulation protocols each | Bootstrap | After stimulation onset<br>95% confidence Interval =<br>0.26-0.30, $p < 0.0001$ (N→W); 0.03-0.04, $p < 0.0001$ (W→N) |
| Fig. 1g | 8 mice, 2 stimulation protocols each | Bootstrap | After stimulation termination<br>95% confidence Interval =<br>0.03-0.04, $p < 0.0001$ (N→W); 0.23-0.28, $p < 0.0001$ (W→N) |
| Fig. 3c (Left) | 6 mice | Bootstrap | After stimulation onset (Left)<br>95% confidence Interval =<br>0.31-0.37, $p < 0.0001$ (N→W); 0.04-0.05, $p < 0.0001$ (W→N) |
| Fig. 3c (Right) | 6 mice | Bootstrap | After stimulation termination (Right)<br>95% confidence Interval =<br>0.016-0.024, $p < 0.0001$ (N→W); 0.027-0.035, $p < 0.0001$ (W→N) |
| Fig. 3j (Top) | 10-19 sessions from 6-7 mice | One-sided Wilcoxon rank-sum test | $W = 494$, $p < 0.0001$ (GCaMP vs. GRAB<sub>NE</sub> in LC),<br>$W = 380$, $p < 0.0001$ (GCaMP8m vs. GRAB<sub>NE</sub> in mPFC) |
| Fig. 3j (Bottom) | 10-15 sessions from 5-6 mice | One-sided Wilcoxon rank-sum test | $W = 270$, $p = 0.0002$ (GCaMP vs. GRAB<sub>NE</sub> in LC),<br>$W = 276$, $p = 0.0037$ (GCaMP8m vs. GRAB<sub>NE</sub> in mPFC) |
| Ext. Data Fig. 3d,e | 5 mice | One-sided Wilcoxon signed-rank test | $W = 15$, $p = 0.03$ (NREM-to-wake)<br>$W = 4$, $p = 0.22$ (wake-to-NREM) |
| Ext. Data Fig. 3f,g | 5 mice | One-sided Wilcoxon signed-rank test | $W = 0$; $p = 0.03$ (NREM-to-wake)<br>$W = 14$, $p = 0.06$ (wake-to-NREM) |

